# A robust approach for preserving and sectioning fragile 3D spheroids for high-quality histological analysis

**DOI:** 10.64898/2026.08.05.743094

**Authors:** Ramón Cervantes-Rivera, Sandra Jetsamari Figueroa Ortíz, Atalia Ziret Romero Rosas, Adrián Sánchez Orozco, Ma. Antonia Herrera-Vargas, Esperanza Meléndez-Herrera, Manuel López Rodríguez, Alejandra Ochoa-Zarzosa, Joel E. López-Meza

## Abstract

Three-dimensional (3D) spheroid models have become essential in cancer biology, drug screening, and tissue engineering. However, their small size, fragile structure, and tendency to disintegrate during routine histoprocessing present persistent technical challenges. Conventional paraffin embedding often results in tissue fragmentation, loss of spatial orientation, and poor section quality, whereas cryosectioning often compromises cellular morphology. Here, we present a robust, cost-effective protocol for preserving and sectioning fragile 3D spheroids, resulting in high-quality histological sections with intact architecture and excellent cellular detail. The method involves optimized handling and embedding procedures that stabilize spheroids during standard formalin fixation, paraffin infiltration, and microtomy, eliminating mechanical distortion and preserving spherical integrity for consistent sectioning. We demonstrate successful application across different cell line spheroids, with subsequent compatibility with hematoxylin and eosin (H&E) staining protocols. Compared to conventional methods, our approach significantly reduces sample loss, improves inter-section reproducibility, and preserves fine structural features such as necrotic cores, proliferative zones, and extracellular matrix components. This protocol provides a reliable, accessible solution for routine histological analysis of fragile 3D spheroids, facilitating more accurate morphological and molecular assessment in translational research settings.

**Key features:**

- Maintains spheroid integrity: Prevents mechanical distortion, fragmentation, and loss of spatial orientation during processing.
- Significantly reduces sample loss: Decreases failure rate compared to traditional methods, conserving valuable samples.
- Broad spheroid compatibility: Works effectively with primary tumor-derived, stem cell-derived, and co-culture spheroid models.
- Enables high-quality sectioning and staining: Delivers consistent, reproducible sections that are fully compatible with H&E, IHC, and IF.

**Graphical overview:** 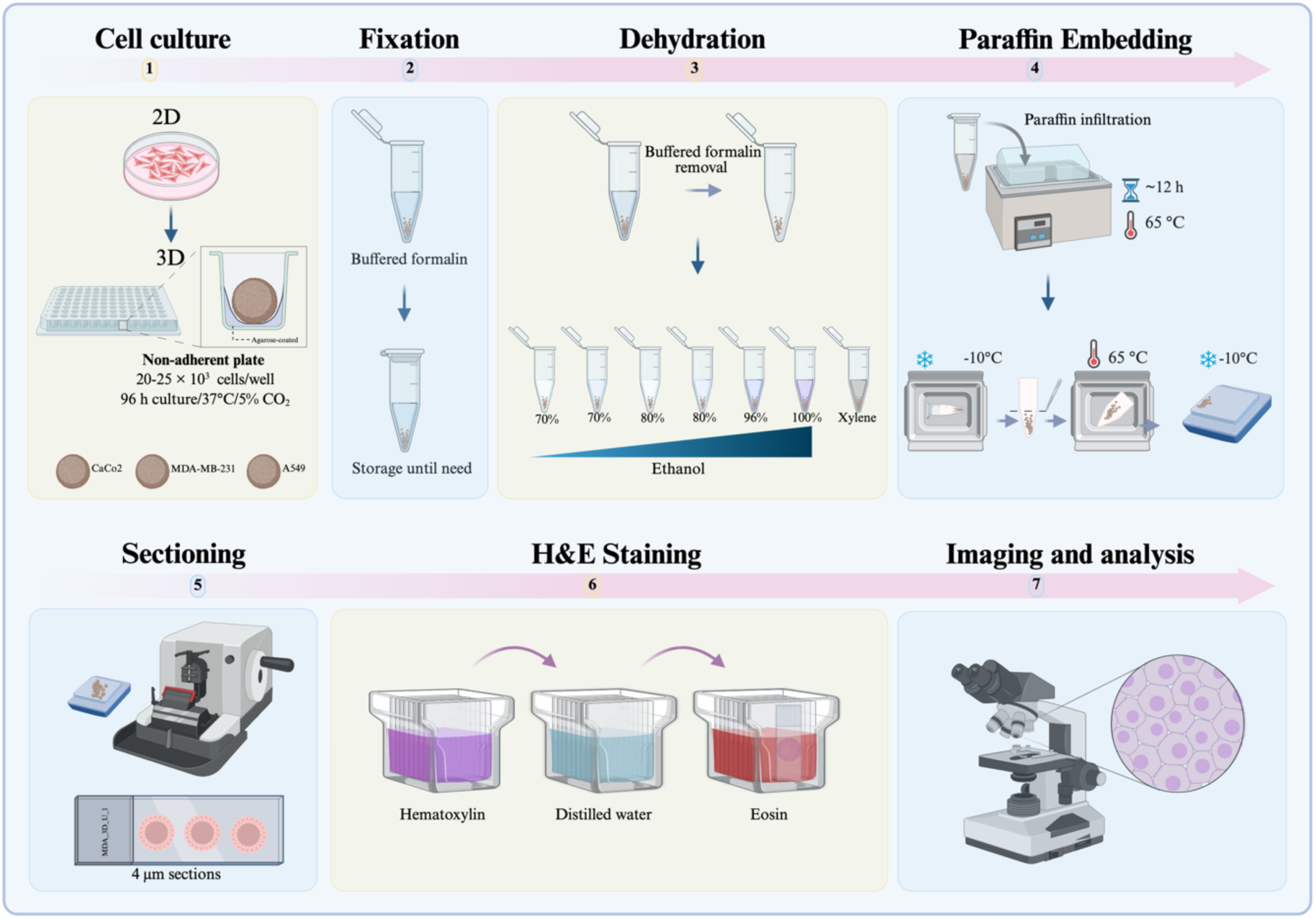

## Background

Three-dimensional (3D) tumor models have advanced to become *in vitro* platforms that more accurately mimic the biological complexity of cancer than traditional two-dimensional (2D) monolayer cultures[1]. Cancer is a multifactorial and diverse disease characterized by uncontrolled cell growth, genomic instability, metabolic reprogramming, immune evasion, and the ability to invade and metastasize[2]. Notably, these processes are heavily influenced by the tumor microenvironment (TME), which includes oxygen and nutrient gradients, extracellular matrix (ECM) composition, biomechanical forces, and cell-to-cell communication[3]. While 2D systems are convenient for experiments, they do not replicate the spatial organization and microenvironmental conditions that control tumor behavior. Cells in 2D cultures assume an unnatural flat shape, with unlimited apical diffusion and limited basal attachment, lacking proper cell-cell contacts and a three-dimensional structure that regulates cell signaling, gene expression, and drug response. As a result, 2D models have limited relevance and predictive ability in preclinical drug testing[1].

3D tumor models include various platforms such as multicellular spheroids, organoids, scaffold-based cultures, and bioprinted constructs. Among these, tumor spheroids are particularly popular and cost-effective[3]. In non-adherent culture conditions, tumor cells spontaneously form compact multicellular aggregates that develop inherent structural organization and physiologically relevant gradients of oxygen, nutrients, metabolites, and signaling molecules. These gradients create spatially distinct cellular zones, including an outer proliferative layer, intermediate quiescent regions, and a central hypoxic or necrotic core, closely mimicking avascular tumor regions *in vivo*[4]. The radial structure of spheroids, with a high-oxygen, nutrient-rich outer layer and a progressively hypoxic, nutrient-deprived center, reflects the key mass-transfer limitations seen in solid tumors. This stratification enables researchers to investigate pathophysiological processes such as oxygen-and nutrient-driven metabolic adaptation, hypoxia-inducible factor (HIF)-mediated signaling, and the transition from quiescence to necrosis in a precisely controlled environment [5].

Establishing spatial heterogeneity within spheroids has significant implications for cancer research. Cells in different regions of the spheroid exhibit distinct transcriptional programs, metabolic states, proliferative capacities, and responses to stress. Hypoxic cores activate pathways such as HIF-mediated signaling, which promote metabolic adaptation and survival under nutrient deprivation [5]. Additionally, increased cell-cell adhesion, changes in polarity, and endogenous ECM deposition create diffusion barriers that affect drug penetration and therapeutic response. Consequently, spheroid models clinically relevant phenomena such as chemoresistance, microenvironment-driven plasticity, and treatment-induced selective pressures more effectively than 2D cultures [6]. Recent studies have demonstrated that including stromal or immune cells -such as co-culturing cancer cells with macrophages-can be integrated into spheroid models to imitate the complex cellular interactions of the TME, allowing tracking of region-specific changes in hypoxia, apoptosis, and metabolic profiles across the radial axis[3, 7].

The use of 3D models greatly influences the study of cancer progression and drug development. Drug screening in spheroids often shows lower drug sensitivity compared to 2D cultures, emphasizing the importance of the microenvironment in treatment response[1]. Additionally, 3D systems allow investigation of tumor-matrix interactions, invasive behavior, and ECM remodeling when combined with biomimetic matrices such as collagen, matrigel, or synthetic hydrogels. These matrix-embedded systems mimic biomechanical constraints and stromal interactions found *in vivo*, offering a more realistic framework for studying tumor adaptation and growth[7]. For example, embedding spheroids in collagen I hydrogels enables direct observation of cells escaping from the tumor mass and invading surrounding tissue, a process key to metastasis and drug resistance[7]. The ability to observe such events in both space and time makes 3D spheroids especially useful for mechanistic research and developing anti-invasive treatments[3].

Despite these advantages, 3D spheroid models pose technical challenges, particularly during downstream histological processing. Because they are small, have compact cellular structures, and lack intrinsic support, spheroids are very prone to deformation, fragmentation, or loss during fixation, dehydration, paraffin embedding, and microtome sectioning [4, 8]. Matrix-embedded spheroids face extra difficulties, as mechanical mismatch or uneven shrinkage between the spheroid and the surrounding hydrogel can weaken their structure [7]. Their small size also makes spheroids difficult to locate and position within a paraffin block, and incomplete dehydration or poor paraffin infiltration can lead to voids, tearing, or complete loss during sectioning [9, 10]. As a result, many researchers choose low-resolution whole-mount imaging or cryosectioning, both of which trade off either spatial detail or molecular preservation [8].

Histological sectioning is crucial for thoroughly characterizing spheroids. While live imaging and viability assays offer overall functional insights, they cannot reveal internal structure or localized treatment effects[11]. Histological analysis allows direct visualization of proliferative zones, hypoxic regions, necrotic cores, and area-specific therapeutic damage at a microscopic level[12]. Additionally, formalin-fixed, paraffin-embedded (FFPE) sections are preferable to whole-mount preparations for many applications, such as immunohistochemistry, because they prevent the depth-related antibody penetration issues seen in thick samples and enable long-term storage in biobanks[8]. Consequently, developing optimized and consistent histological workflows is essential for unlocking the full analytical potential of 3D tumor spheroid systems[9, 10].

In this context, developing tailored, step-by-step histological protocols specifically adapted for 3D spheroid models is necessary to ensure structural preservation, reproducibility, and reliable spatial analysis[4, 13]. These methodological improvements support the wider adoption of 3D tumor models in cancer biology research, drug discovery, and preclinical evaluation[1, 3].

## Materials and reagents

### Biological materials

1. MDA-MB-231 cell line (ATCC, catalog number: CRM-HTB-26)
2. A549 cell line (ATCC, catalog number: CRM-CCL-185)
3. Caco-2 cell line (ATCC, catalog number: HTB-37)
4. Three-dimensional (3D) spheroids of all cell lines (see Recipes)

### Reagents

1. 10% Neutral buffered formalin (J.T. Baker, catalog number: 2106-03)
2. Phosphate-buffered saline (1**×** PBS) (commercial or lab-made)
3. Ethanol (Meyer, catalog number: 0390)
4. Xylene (J.T. Baker, catalog number: 9490-03)
5. Paraffin wax, histology grade (melting point 56-60 °C) (Sigma-Aldrich, catalog number: 1.07164.2504)
6. Hematoxylin solution (Hycel, catalog number: 738)
7. Eosin Y solution (Sigma-Aldrich, catalog number: 45380)
8. Lithium carbonate (Fagan Lab, catalog number: 2518)
9. Synthetic resin for mounting (Sigma-Aldrich, catalog number: 1.07960.0500)
10. Methanol (J.T. Baker, catalog number: 9070-03)
11. Dulbecco’s modified Eagle medium (DMEM) (Sigma Merck, catalog number: D5648) or RPMI media.
12. Sodium bicarbonate (NaHCO_3_) (J.T Baker, catalog number: 3506-01)
13. L-glutamine (Gibco, catalog number: 20530-081)
14. Amphotericin B (Sigma-Aldrich, catalog number: A2942)
15. Fetal bovine serum (FBS, BioWest, catalog number: BIO-S1400)
16. Bovine calf serum (BCS, BioWest, catalog number: S0400-500)
17. Penicillin-streptomycin (Gibco, catalog number: 15140-122)
18. Trypsin (Sigma-Aldrich, catalog number: T4799-5G)
19. Ethylenediaminetetraacetic acid (EDTA) (J.T Baker, catalog number: 8993-01)
20. Dimethyl sulfoxide (DMSO) (Sigma-Aldrich, catalog number: D2650)
21. Actinomycin D (Sigma-Aldrich, catalog number: A9415)
22. Potassium chloride (J.T. Baker, catalog number: 3040-01)
23. Sodium chloride (J.T. Baker, catalog number: 3624-01)
24. Disodium phosphate (Na_2_HPO_4_) (J.T Baker, catalog number: 3828-01)
25. Monobasic potassium phosphate (KH_2_PO_4_) (J.T Baker, catalog number: 3246-01)
26. Agarose (Invitrogen, catalog number: 16500-100)
27. MilliQ water sterile water (J.T. Baker, catalog number: 4220-20)
28. Hydrochloric acid (HCl) (J.T. Baker, catalog number: 9535-05)
29. Sodium hydroxide (NaOH) (Macron, fine chemicals, catalog number: 7708-10)

### Laboratory supplies

1. 96-well cell culture plate (NEST, catalog number: 701002)
2. Cell culture dish (60 mm × 15 mm) (NEST, catalog number: 705007)
3. Conical-bottom centrifuge tube, 50 mL (Uniparts, catalog number: 32117F)
4. Conical-bottom centrifuge tube, 15 mL (Uniparts, catalog number: 34117F)
5. Microcentrifuge tubes 1.5 mL (Axygen, catalog number: MCT-150-C)
6. Pipette tip 100-1000 µL (Uniparts, catalog number: 51131)
7. Pipette tip 1-200 µL (Uniparts, catalog number: 51121Y)
8. Pipette tip 0.5-10 µL (Axygen, catalog number: T-300)
9. Embedding molds (CR Globe, catalog number: P2030)

### Solutions

a. Culture medium for MDA-MB-231, Caco-2, and A-549 cell lines (DMEM complete medium) (see Recipes)
b. 1.5% agarose solution (see Recipes)
c. 1× PBS solution (see Recipes)
d. 0.04% Trypsin-EDTA (see Recipes)
e. 500 mM Actinomycin D (see Recipes)
f. 70%, 80%, 96% and 100% ethanol solutions (see Recipes)
g. Eosin solution (see Recipes)
h. 10 % neutral buffered formalin (see Recipes)
i. Lithium carbonate bluing solution (see Recipes)

## Recipes

### a) Culture medium for MDA-MB-231 cells: Dulbecco’s Modified Eagle Medium (DMEM-complete)

a) Culture medium for MDA-MB-231 cells: Dulbecco's Modified Eagle Medium (DMEM-complete)
| Reagent | Final concentration | Quantity |
| --- | --- | --- |
| DMEM powder | - | As required |
| Sodium bicarbonate (NaHCO<sub>3</sub>) | 3 g/L | 3 g |
| FBS | 5% (v/v) | 50 mL |
| BCS | 5% (v/v) | 50 mL |
| Penicillin-streptomycin | 1% (v/v) | 10 mL |
| Amphotericin B | 0.05% (v/v) | 500 µL |
| L-glutamine (200 mM) | 0.05% (v/v) | 500 µL |
| Sterile distilled water | - | To 1 L final volume |

Cells base medium preparation:

1. Combine the DMEM powder with sterile distilled water and stir continuously until fully dissolved.
2. Add 3 g of sodium bicarbonate (NaHCO₃) to the solution.
3. Adjust the pH to 7.4 by carefully adding 1 M NaOH or 1 M HCl dropwise as required.

Supplementation (perform inside a laminar flow hood):

- Add the following components to the base medium:

1. 5% fetal bovine serum (FBS) → 50 mL
2. 5% bovine calf serum (BCS) → 50 mL
3. 1% penicillin–streptomycin → 10 mL
4. 0.05% amphotericin B → 500 µL
5. 0.05% L-glutamine (200 mM stock) → 500 µL

Final Sterilization and storage:

- Filter-sterilize the complete medium using a vacuum filtration system equipped with a 0.22 µm MCE membrane.
- Store the sterile medium at 4 °C and use within 4 weeks.

**Note 1:** This formulation is routinely used for culturing MDA-MB-231 cells, and RPMI medium works.

**Note 2:** The formulation for Caco-2 and A549 is the same, but without 0.05% L-glutamine.

### b) 1.5 % agarose solution

| Reagent | Final concentration | Quantity |
| --- | --- | --- |
| Agarose | 1.5% (w/v) | 1.5 g |
| Triple-distilled water | - | To 100 mL |
| Final volume | - | 100 mL |

1. In a glass flask, combine 1.5 g of agarose with 100 mL of triple-distilled water.
2. Autoclave the mixture for 15 minutes to achieve sterility.

**Note:** This solution is intended to create non-adherent surfaces in standard 96-well plates. After autoclaving, it should be used immediately or kept at approximately 60-70 °C until ready to dispense.

### c) 1× PBS (pH 7.4)

| Reagent | Final concentration | Quantity |
| --- | --- | --- |
| NaCl | 137 mM | 8.0 g |
| KCl | 2.7 mM | 0.2 g |
| Na<sub>2</sub>HPO<sub>4</sub> | 10 mM | 1.44 g |
| KH<sub>2</sub>PO<sub>4</sub> | 1.8 mM | 0.24 g |
| ddH<sub>2</sub>O | - | 1 L |

1. Dissolve NaCl (8.0 g), KCl (0.2 g), Na₂HPO₄ (1.44 g), and KH₂PO₄ (0.24 g) in approximately 800 mL of distilled water using a magnetic stirrer. Continue stirring until the solution is completely clear and free of any undissolved particulates.
2. Calibrate the pH meter using standard buffer solutions. Then, while continuously stirring, titrate the solution to pH 7.4 by adding 1 M HCl or 1 M NaOH drop by drop. Allow the pH reading to stabilize for 15-20 seconds after each addition before recording the value.
3. Quantitatively transfer the solution to a 1 L volumetric flask and bring the total volume to exactly 1 L with distilled water. Mix thoroughly by inverting the capped flask at least five times to ensure homogeneity.
4. Pour the solution into an autoclave-safe bottle, loosen the cap slightly to permit pressure equilibration, and autoclave at 121 °C (15 psi) for 20-30 minutes.
5. After autoclaving, let the solution cool to room temperature before use. If not used immediately, store it under appropriate conditions (e.g., 4 °C for short-term storage) and label the bottle with the solution name, pH, and preparation date.

### d) 0.04 % Trypsin-EDTA

| Reagent | Final concentration | Quantity |
| --- | --- | --- |
| Trypsin | 0.04% (w/v) | 390 mg |
| EDTA | 0.037% (w/v) | 370 mg |
| Sodium chloride (NaCl) | - | 85 mg |
| PBS 1× | - | To 1 L |
| Final volume | - | 1 L |

1. Use 1 L of 1× PBS (homemade or commercial).
2. In a separate sterile container, dissolve 390 mg of trypsin and 85 mg of NaCl in 10 mL of the prepared PBS. Mix gently until completely dissolved.
3. Add 370 mg EDTA to the remaining PBS (about 990 mL) and stir until fully dissolved.
4. Add the trypsin-NaCl solution to the EDTA-containing PBS and stir thoroughly to ensure even mixing.
5. Under aseptic conditions, filter-sterilize the combined solution using a 0.22 µm vacuum filtration unit.
6. Divide the sterile solution into appropriate aliquots, clearly label each with the concentration and date, and store at -20 °C until use.

### e) 500 µM actinomycin D stock solution

| Reagent | Final concentration | Quantity |
| --- | --- | --- |
| Actinomycin D | 500 µM | As required |
| DMSO | - | As required |

Actinomycin D is a powerful cytotoxic and teratogenic agent. Always wear appropriate personal protective equipment (PPE), such as gloves and safety goggles, and handle all materials in a designated chemical fume hood or biological safety cabinet.

1. Under sterile conditions in a cell culture hood, dissolve the required amount of Actinomycin D in sterile DMSO to a final concentration of 500 µM.
2. Gently mix by pipetting or vortexing until the compound is fully dissolved.
3. Transfer the solution into amber microcentrifuge tubes or tubes wrapped in aluminum foil to protect from light.
4. Clearly label each aliquot with the compound name, concentration, and preparation date.
5. Store the aliquots at -20 °C, protected from light. Avoid repeated freezing and thawing.

### f) Ethanol solutions

| Reagent | Final concentration | Quantity or Volume |
| --- | --- | --- |
| **Absolute ethanol** | 70/80/96% (v/v) | 70/80/96 mL |
| **Distilled water** | - | 30/20/4 mL |
| **Final volume** | - | 100 mL |

To prepare 100 mL of each concentration, mix absolute ethanol with distilled water as follows:

- 70% ethanol: 70 mL absolute ethanol + 30 mL distilled water
- 80% ethanol: 80 mL absolute ethanol + 20 mL distilled water
- 96% ethanol: 96 mL absolute ethanol + 4 mL distilled water Combine the two components in a suitable container and mix thoroughly.

### g) Eosin solution

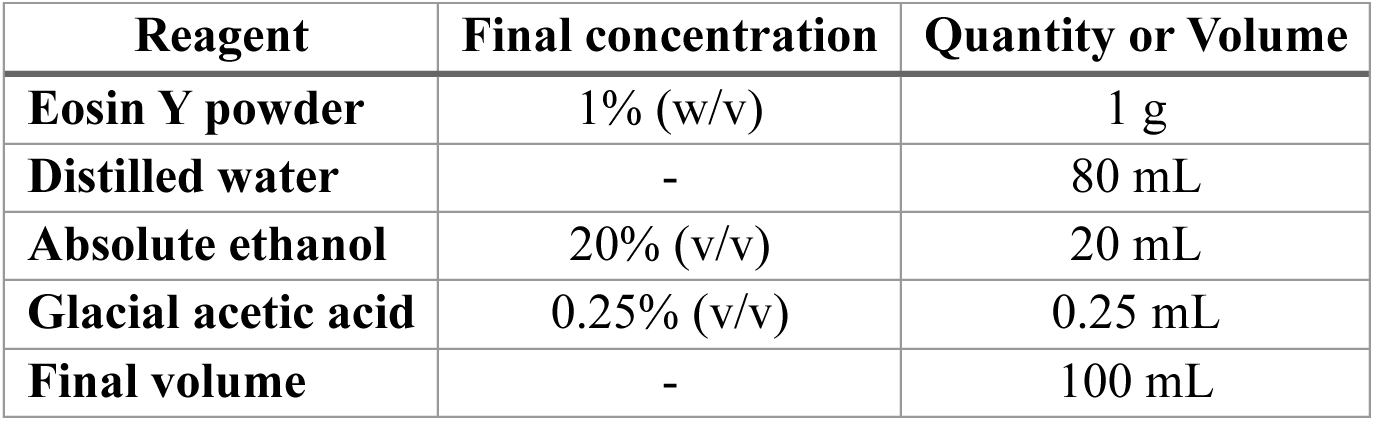

| Reagent | Final concentration | Quantity or Volume |
| --- | --- | --- |
| **Eosin Y powder** | 1% (w/v) | 1 g |
| **Distilled water** | - | 80 mL |
| **Absolute ethanol** | 20% (v/v) | 20 mL |
| **Glacial acetic acid** | 0.25% (v/v) | 0.25 mL |
| **Final volume** | - | 100 mL |

1. In a glass container, dissolve 1 g of eosin Y powder in 80 mL of distilled water.
2. Add 20 mL of absolute ethanol and mix until the powder is fully dissolved.
3. Add 0.25 mL of glacial acetic acid to enhance cytoplasmic staining.
4. Carefully stir the solution until it’s uniform.
5. If necessary, filter the solution, then transfer it to an amber bottle and store it at room temperature.

### h) 10% neutral buffered formalin (from 37% stock)

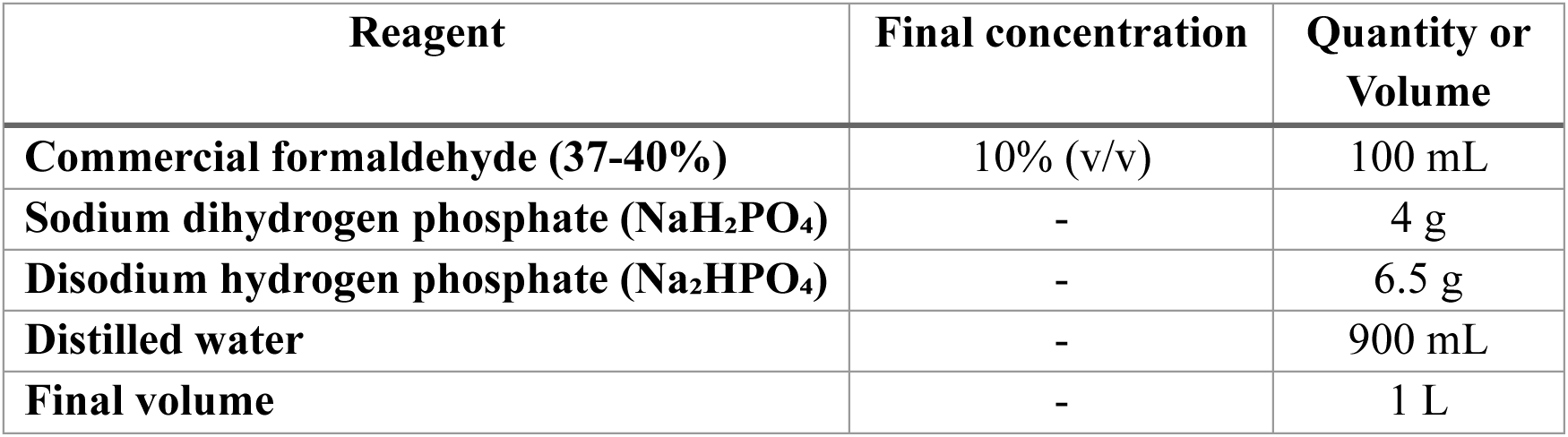

| Reagent | Final concentration | Quantity or Volume |
| --- | --- | --- |
| **Commercial formaldehyde (37-40%)** | 10% (v/v) | 100 mL |
| **Sodium dihydrogen phosphate (NaH<sub>2</sub>PO<sub>4</sub>)** | - | 4 g |
| **Disodium hydrogen phosphate (Na<sub>2</sub>HPO<sub>4</sub>)** | - | 6.5 g |
| **Distilled water** | - | 900 mL |
| **Final volume** | - | 1 L |

1. Dissolve 4 g of sodium dihydrogen phosphate (NaH₂PO₄) and 6.5 g of disodium hydrogen phosphate (Na₂HPO₄) in 900 mL of distilled water.
2. Add 100 mL of commercial formaldehyde solution (37-40%).
3. Mix thoroughly and confirm that the final pH is approximately 7.0.
4. Label the container and store at room temperature.

### i) Lithium carbonate bluing solution

| Reagent | Final concentration | Quantity or Volume |
| --- | --- | --- |
| Lithium carbonate ($\text{Li}_2\text{CO}_3$) | 0.1% (w/v) | 0.1 g |
| Distilled water | - | 100 mL |
| Final volume | - | 100 mL |

1. Weigh 0.1 g of lithium carbonate and dissolve it in 100 mL of distilled water.
2. Mix thoroughly until fully dissolved.
3. Label the container.
4. Store at room temperature. For best results, prepare fresh when possible.

### j) 1 M HCl

| Reagent | Final concentration | Quantity |
| --- | --- | --- |
| 37% HCl | 1 M | 9.05 mL |
| MilliQ water | - | Up to 100 mL |

1. Put on gloves and safety goggles. Work inside a fume hood.
2. Measure 9.05 mL of 37% HCl using a graduated cylinder or pipette.
3. Add approximately 50-70 mL of Milli-Q water to a 100 mL volumetric flask.
4. Slowly pour the measured HCl into the water while gently swirling the flask.
5. Allow the solution to cool to room temperature (the flask will become warm).
6. Top up with Milli-Q water to the 100 mL mark (meniscus at eye level).
7. Mix thoroughly by inverting the capped flask at least five times.
8. Transfer to a labeled glass bottle and store at room temperature.

**Caution:** Always add acid to water, never water to acid.

### k) 1 M NaOH

| Reagent | Final concentration | Quantity |
| --- | --- | --- |
| NaOH | 1 M | 4 g |
| MilliQ water | - | Up to 100 mL |

1. Put on gloves and safety goggles. Work in a well-ventilated area or inside a fume hood.
2. Weigh 4.0 g of NaOH pellets using an analytical balance.
3. Add approximately 50-70 mL of Milli-Q water to a 100 mL volumetric flask (or a glass beaker for initial dissolution).
4. Add the NaOH pellets slowly to the water while gently stirring with a glass rod.
5. Allow the solution to cool to room temperature (the dissolution is exothermic).
6. If using a beaker, quantitatively transfer the solution to the volumetric flask. Bring the final volume to the 100 mL mark with Milli-Q water (meniscus at eye level).
7. Mix thoroughly by inverting the capped flask at least five times.
8. Transfer to a labeled plastic or glass bottle and store at room temperature.

**Caution:** NaOH is caustic. Always add NaOH to water, not water to NaOH, to avoid violent boiling and splashing.

## Equipment

1. Centrifuge (PowerSpin™, model: C856)
2. Micropipettes (Axygen, AP-10, AP-100 and AP-1000):

a. 0.5-10 µL Single-channel Pipettor (Axygen® Axypet®, catalog number: AP-10)
b. 10-100 µL Single-channel Pipettor (Axygen® Axypet®, catalog number: AP-100)
c. 100-1,000 µL Single-channel Pipettor (Axygen® Axypet®, catalog number: AP-1000)
3. Orbital shaker (Benchmark, model: BT302)
4. pH meter (Apera, PH700)
5. Digital stirring hot plate (Thermo Scientific, model: SP131015Q)
6. Inverted microscope (Carl Zeiss, model: 37081)
7. Tissue flotation bath (Chicago Surgical & Electrical Co., Cat. No. 26103)
8. Tissue embedding center (Ecoshel, model: ECO-6L)
9. Rotary microtome (Microm International, model: HM325)
10. Hot plate (Thermolyne, model: type 2200)
11. Spin tissue processor (Microm International, model: STP-120)
12. Laminar flow cabinet (Thermo Fisher Scientific, model: 1340)
13. CO₂ incubator for cell culture (Thermo Fisher Scientific, model: 3422)
14. Neubauer chamber (Marienfeld, catalog number: 0610010)

## Procedure

### I. Preparation of cost-effective non-adherent plates using agarose coating

1. **Prepare the agarose solution**. Make a sterile 1.5% (w/v) agarose solution (see Recipes). Keep it molten by placing the container in a 90 °C water bath on a hot plate with stirring to prevent gelling.
2. Use a sterile, flat-bottom 96-well cell culture plate as the substrate for coating.
3. **Prepare a working aliquot**. Under aseptic conditions, transfer 1 mL of molten agarose to a sterile 1.5 mL microcentrifuge tube. This aliquot will help prevent contamination of the main stock.
4. **Coat the wells**. Using the working aliquot, immediately dispense 60 µL of molten agarose into each well of the 96-well plate (Figure 1).

**Note:** Avoid using a multichannel pipette because standard reagent reservoirs cannot withstand the high temperature needed to keep agarose molten. Work quickly with a single-channel pipette to prevent the agarose from solidifying inside the tip or tube.

1. **Refresh the aliquot if necessary**. Repeat the dispensing step until all required wells are filled. If the agarose in the working aliquot begins to solidify, replace it with fresh molten agarose from the main stock.
2. **Solidify and dry**. Allow the agarose to solidify undisturbed at room temperature. Once solid, leave the plate uncovered in a cell culture hood for 15 minutes to allow residual moisture to evaporate under laminar flow.
3. **Seal the plate**. Wrap it tightly with clean plastic film (e.g., Parafilm) and place it inside a sealed plastic bag to prevent dehydration and contamination.
4. **Store**. Keep the coated plates at 4 °C. Under these storage conditions, they remain usable for 2-3 weeks.

**Figure 1.**
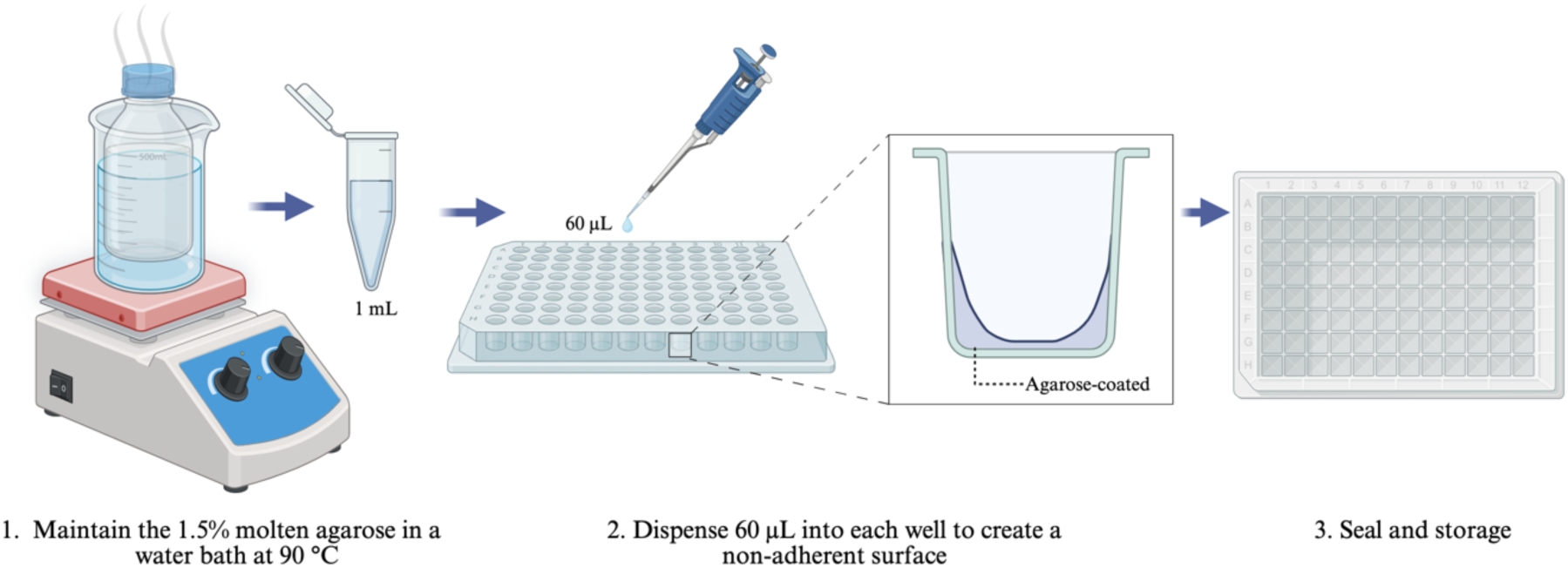
Preparation of cost-effective, non-adherent plates using agarose coating.

### II. Thawing MDA-MB-231 cells

1. Remove the cryovial of MDA-MB-231 cells from liquid nitrogen.
2. Warm it immediately in a 37°C water bath, swirling gently until completely thawed (about 1-2 minutes).
3. Inside a biosafety cabinet, use a sterile 2 mL serological pipette to transfer the thawed cells into a sterile 1.5 mL tube.
4. Centrifuge the tube at 257 × g for 5 minutes at room temperature to form a cell pellet.
5. Carefully remove the supernatant with a sterile 2 mL serological pipette, avoiding disturbance to the pellet.
6. Gently resuspend the pellet in 1 mL of pre-warmed complete DMEM by pipetting up and down.
7. Transfer the suspension to a 60 mm × 15 mm culture dish, then add 4 mL of complete DMEM.
8. Gently rock the dish in a crosswise pattern to evenly distribute the cells, then place it in a 37°C, 5% CO₂ humidified incubator.

**Note:** If necessary, replace RPMI with DMEM.

### III. Subculture and preparation of MDA-MB-231 Cells

1. Grow MDA-MB-231 cells in 60 mm dishes with complete DMEM at 37 °C, 5% CO₂ until they reach about 80% confluence.
2. Suck off the old medium with a sterile serological pipette.
3. Gently rinse the cells with 5 mL of sterile 1× PBS to remove any remaining serum (which can inhibit trypsin). Then, pour off the PBS.
4. Add 3 mL of warm 0.04% trypsin-EDTA, ensuring it covers the entire cell layer.
5. Let the dish sit at 37 °C for 3-5 minutes. Check under a microscope periodically; cells are ready when they appear round and begin floating.
6. Add 6 mL of complete DMEM (the serum inhibits the trypsin).
7. Gently pipette up and down to break up clumps, then transfer the entire suspension into a sterile 15 mL conical tube.
8. Spin at 257 × g for 5 minutes at room temperature.
9. Carefully pour off the liquid without losing the cell pellet.
10. Add 1 mL of fresh complete DMEM and gently pipette to resuspend the pellet.
11. Count your cells before using them in experiments or for the next passage.

**Note 1:** You can substitute DMEM for RPMI if needed.

**Note 2:** After thawing, pass cells at least 3 times before experiments to allow them to recover and grow steadily.

### IV. Cell counting with a Neubauer chamber (Figure 2)

1. Prepare a 1:10 dilution of your cell suspension:

- Add 90 µL of complete DMEM medium to a sterile 1.5 mL microcentrifuge tube.
- Add 10 µL of the well-mixed cell suspension to the same tube.
- Mix gently by pipetting.
2. Load the hemocytometer:

- Draw 10 µL of the diluted suspension into the chamber of a Neubauer hemocytometer. Allow the liquid to fill the chamber by capillary action. The chamber should be fully covered but not overflowing.
3. Count the cells:

- Place the hemocytometer under an inverted microscope.
- Count the cells in each of the four corner squares (each contains 16 small squares). **Rule:** Include cells touching the top and left edges. Exclude cells touching the bottom and right edges.
4. Calculate the concentration (cells/mL):

- Average the counts from the four corner squares.
- Multiply by 10,000 (hemocytometer conversion factor).
- Multiply by 10 (dilution factor).

Formula:

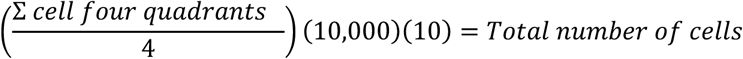

**Figure 2.**
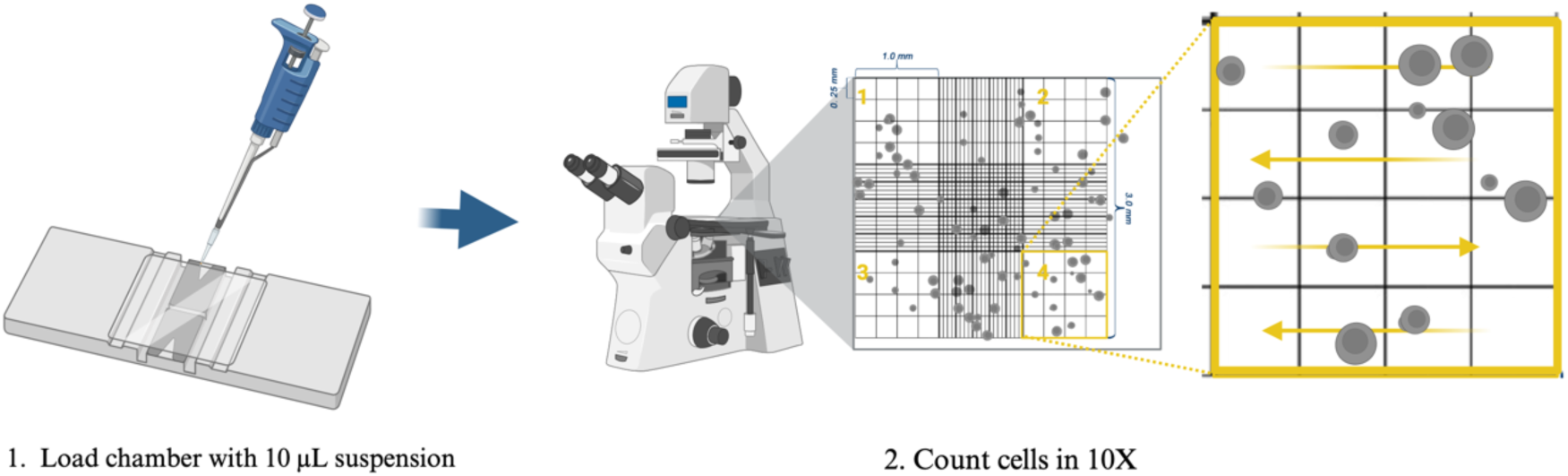
Cell counting with the Neubauer chamber.

Example calculation:

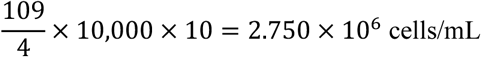

### V. Spheroid formation (**Figure 3**)

1. **Grow cells in a monolayer**. Maintain cells in standard 2D culture conditions (37 °C, 5% CO₂) until they reach the desired confluency.
2. **Prepare the cell suspension**. Resuspend the cells in complete DMEM medium to a final density of 20,000 cells per 100 µL.
3. **Seed the NA plate**. Add 100 µL of the cell suspension to each well of the NA plate.
4. **Promote aggregation**. Place the plate on an orbital shaker set to 150 rpm for 3 hours to encourage cell clumping.
5. **Incubate to form spheroids**. Transfer the plate to a humidified incubator at 37 °C with 5% CO₂, and leave it undisturbed for 96 hours.
6. **Maintain the culture**. Every 48 hours, replace half of the medium in each well, taking care not to disrupt the developing spheroids.
7. **Verify spheroid formation**. Examine the wells under an inverted microscope. Successful formation is indicated by the presence of compact, spherical aggregates.

### VI. Spheroid fixation (**Figure 4**)

1. After spheroid formation is complete, carefully transfer the spheroids into a sterile 1.5 mL microcentrifuge tube using a 1000 µL pipette.
2. Centrifuge at 257 × g for 5 minutes at room temperature to gently pellet the spheroids.
3. Carefully remove the supernatant without disturbing the pellet.
4. Add 50 µL of buffered formalin to completely cover the spheroids.
5. Incubate at room temperature until further downstream processing (at least 24 h).

### VII. Dehydration of fixed spheroids (**Figure 5**)

After fixation, samples should be dehydrated using a graded ethanol series to eliminate water, preparing them for clearing and subsequent embedding. Perform all steps at room temperature.

1. **Replace the fixative with 70% ethanol.** Carefully remove the buffered formalin from the tube containing the fixed spheroids, taking care not to aspirate the pellet. Add 50 µL (or sufficient volume to fully immerse the spheroids) of 70% ethanol. Incubate for 60 minutes at room temperature (repeat twice).
2. **Replace with 80% ethanol.** Remove the 70% ethanol and add 50 µL of 80% ethanol. Incubate for another 60 minutes at room temperature (repeat twice).
3. **Replace with 96% ethanol.** Remove the 80% ethanol and add 50 µL of 96% ethanol. Incubate for a final 60 minutes.
4. **Replace with 100% ethanol.** Remove the 96% ethanol and add 50 µL of 100% ethanol. Incubate for a total of 60 minutes. This step removes most of the remaining water from the samples.
5. **Clear the samples with xylene.** Remove the 100% ethanol and add 50 µL of xylene, or enough to cover the spheroids. Xylene replaces the ethanol, making the spheroids transparent and ready for paraffin infiltration. Incubate at room temperature for 60 minutes. Avoid extending xylene exposure beyond the necessary time to prevent excessive hardening of the sample.

**Figure 3.**
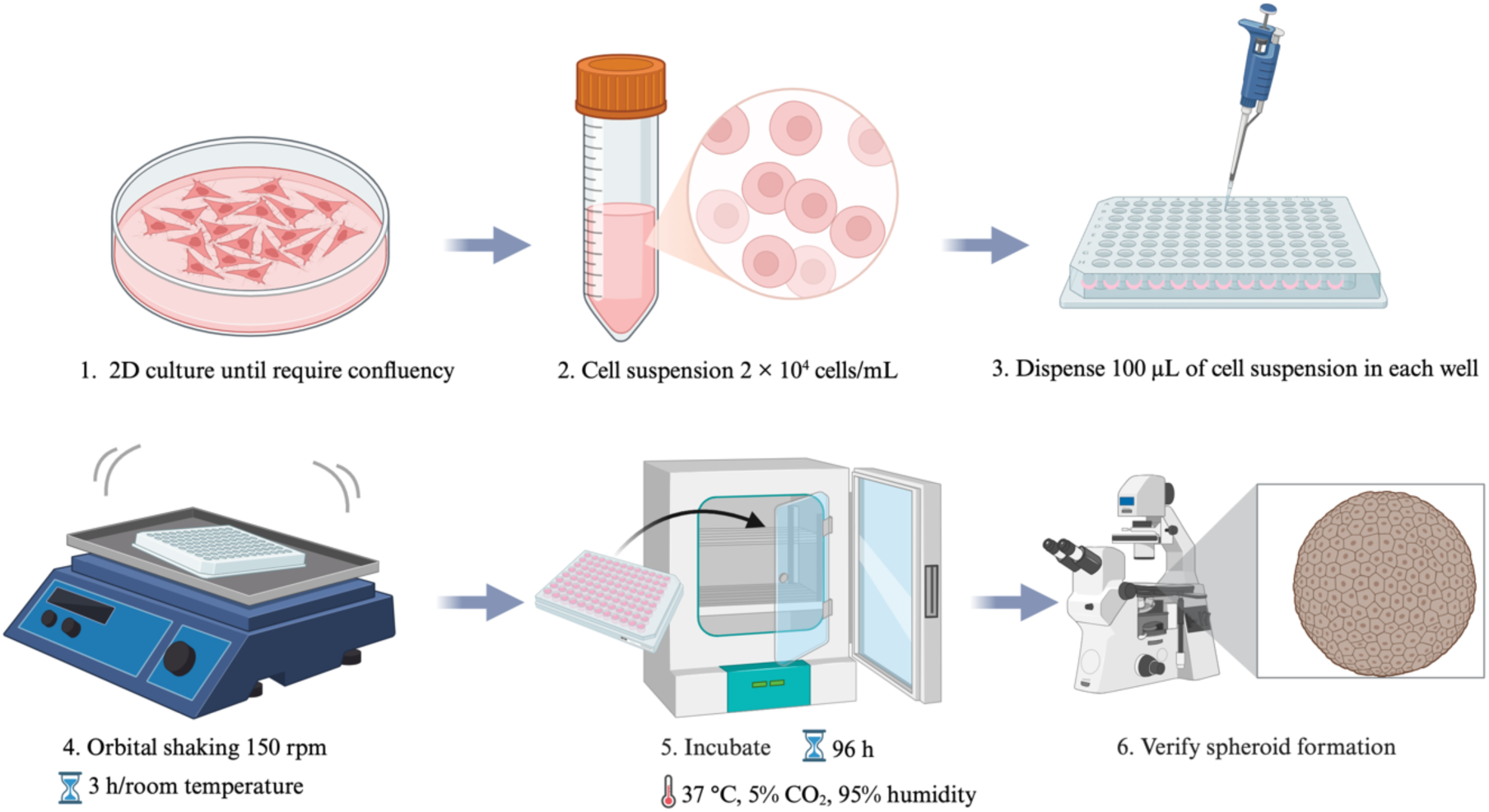
Formation of spheroids in non-adherent plates.

**Figure 4.**
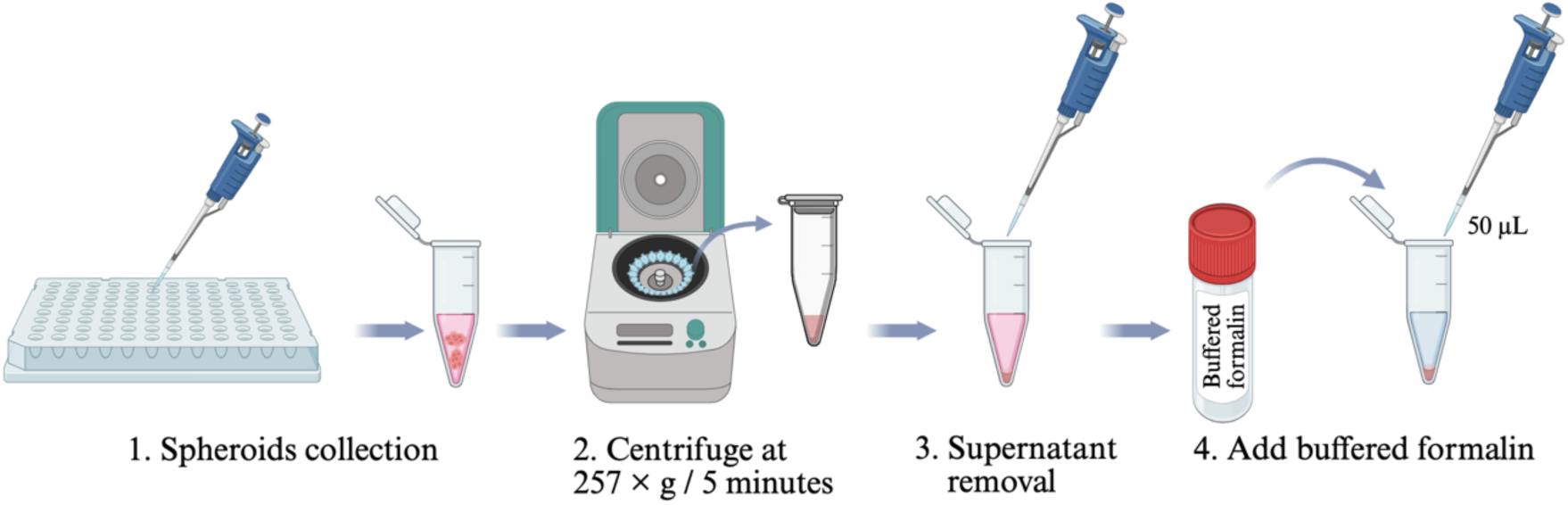
Fixation of spheroids in buffered formalin before downstream processing.

**Figure 5.**
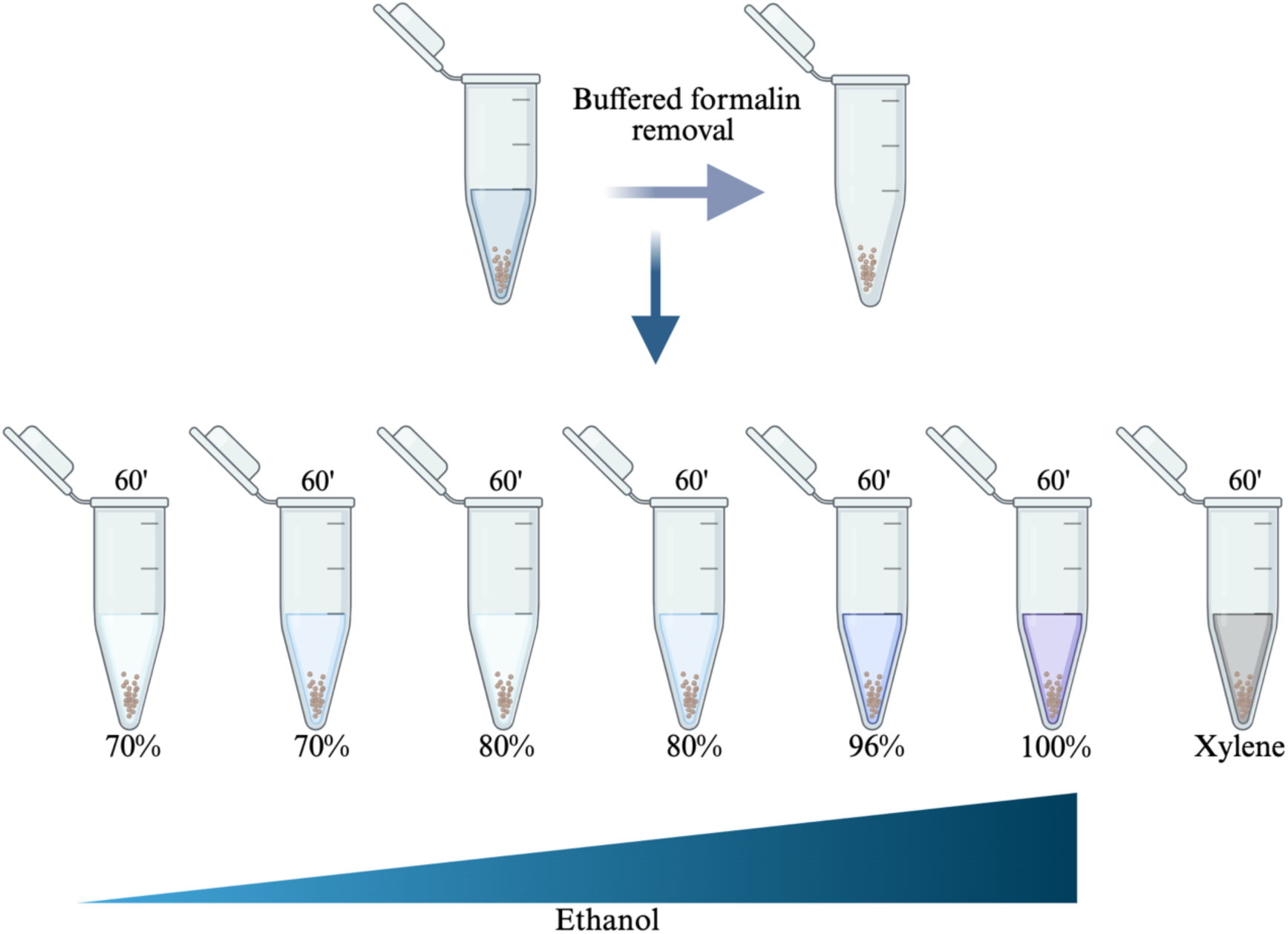
Dehydration and clearing of fixed spheroids for paraffin infiltration.

**Note:** For delicate spheroids, reduce incubation times to 30 minutes per ethanol step to minimize shrinkage. Always perform xylene clearing in a chemical fume hood due to its toxicity.

### VIII. Paraffin infiltration and embedding of samples (**Figure 6**)

After dehydration and clearing, samples must be infiltrated with molten paraffin and embedded in blocks to provide structural support for microtomy. Perform all steps in a well-ventilated area or in a fume hood if using fresh paraffin containing volatile additives. Use a paraffin oven or bath set to exactly 65 °C.

1. **Transfer samples into molten paraffin at 65 °C using a 1.5 mL tube with a floating support.** Spheroids are too small to handle individually; use a floating system to support the 1.5 mL tube during infiltration in the paraffin bath.

- Place the cleared spheroids (from xylene) into a 1.5 mL tube. Add enough molten paraffin at 65 °C to fully cover the spheroids.
- Float the 1.5 mL tube in a 65 °C paraffin bath so that the paraffin inside remains molten.
- Ensure the tube is stable.
2. **Allow infiltration overnight (∼24 h).** Keep the floating 1.5 mL tube in the 65 °C paraffin bath for approximately 24 hours. Do not exceed 65 °C, as higher temperatures may damage the spheroids’ morphology.
3. **Remove samples and place them on a cold plate to make handling easier.** Using pre-warmed forceps, lift the 1.5 mL tube from the paraffin bath. Insert the forceps into the tube, then immediately transfer it to a cold plate (surface temperature -10°C to 4°C). Leave it on the cold plate for 10-20 minutes to let the paraffin solidify. If a cold plate is unavailable, use an ice block.
4. **Transfer to embedding molds with fresh paraffin.**

- Once the paraffin has solidified, carefully lift the forceps to remove the plug. Make sure the bottom part, where the spheroids are concentrated, comes out first. The spheroids will be located at the bottom of the 1.5 mL tube.
- Using a cutter, cut off the bottom portion of the paraffin plug that comes out of the tube; this is where the spheroids are concentrated.
- Place the cut piece into an embedding mold, then add fresh molten paraffin at 65 °C to embed the spheroids.
- Insert the plug into a standard embedding mold filled with fresh molten paraffin at 65 °C, with the spheroid-rich end facing the intended cutting surface.
- Add more fresh molten paraffin to completely fill the mold.
- Use warm forceps or needles to adjust orientation if needed. Always use fresh paraffin for this final embedding to avoid contamination.
5. **Allow blocks to solidify at -10 °C.** Place the filled molds on a level surface in a freezer or on a cold plate set to -10 °C. Solidify for 15-30 minutes, depending on mold size. Rapid cooling to -10 °C produces fine, uniform paraffin crystals, improving sectioning quality. Avoid slow cooling (e.g., at 4 °C), which leads to large crystals that tear sections. After full solidification, remove the paraffin block from the mold-flex plastic molds or briefly warm metal molds.

**Figure 6.**
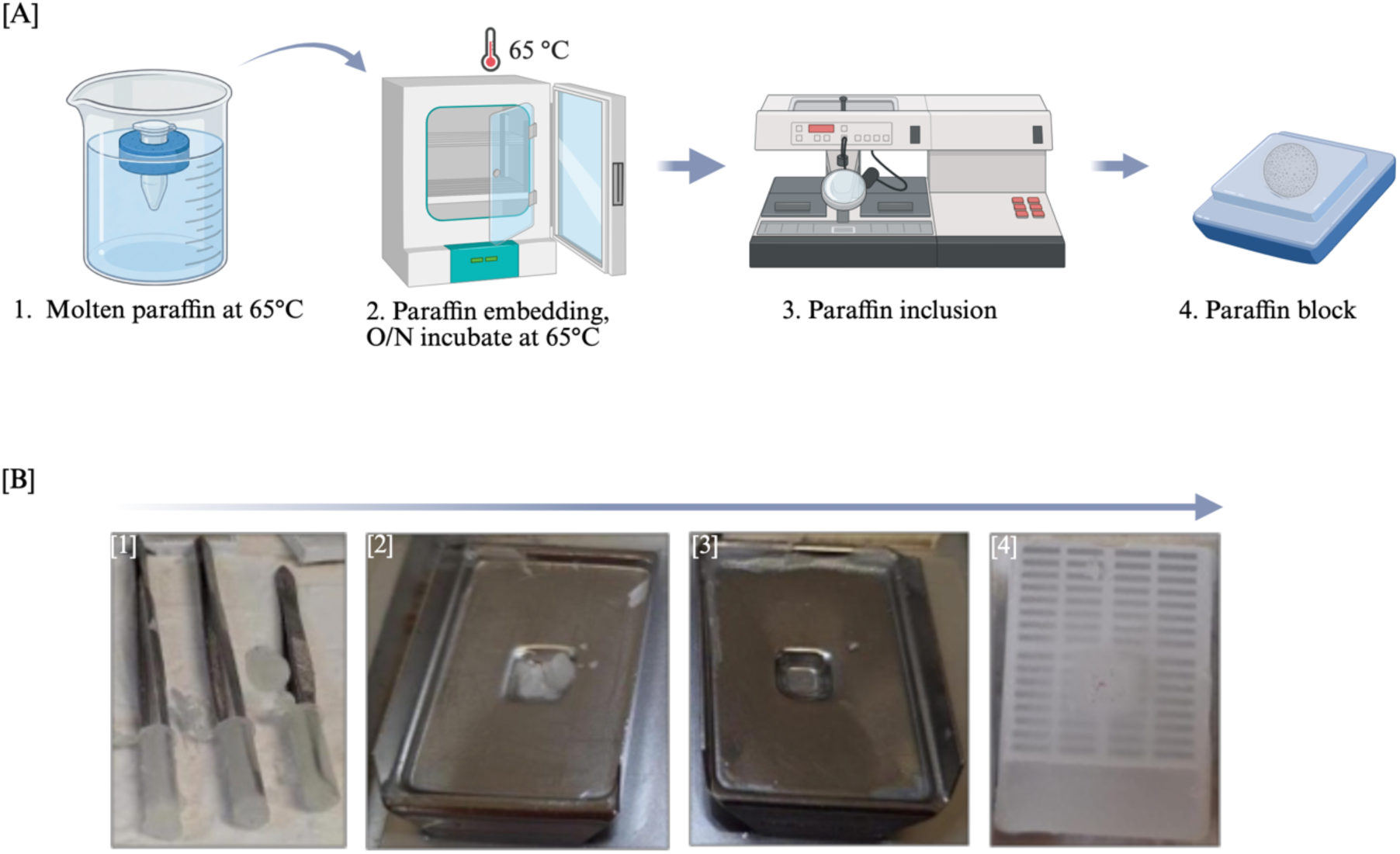
Paraffin embedding of spheroids using a 1.5 mL tube floating system. a) Schematic representation of the paraffin embedding workflow. b) Retrieval of the embedded specimen from a 1.5 mL microtube, using forceps to stabilize the paraffin block.

### IX. Sectioning of paraffin blocks (Figure 7)

After embedding, paraffin blocks must be sectioned to obtain thin ribbons of spheroids for mounting and subsequent staining. Use a rotary microtome and clean glass slides, and perform all steps at room temperature.

1. **Mount the paraffin block in the microtome.** Trim the excess paraffin around the tissue (spheroid-containing area) using a scalpel, leaving a clean block face with the sample fully exposed. Clamp the block securely into the microtome block holder, ensuring the cutting surface is parallel to the blade. Orient the block so that the spheroids will be sectioned at the desired plane, and use a fresh, sharp disposable microtome blade to prevent tearing or compression.
2. **Cut sections at 4 µm thickness.** Set the microtome to cut at 4 µm, advance the block slowly, and discard the first few sections until the full face of the spheroids is exposed. Cut at a steady, moderate speed about one to two sections per second and collect a ribbon of five to ten serial sections. If sections curl or wrinkle, adjust the blade clearance angle. For brittle spheroids, chill the block on ice for five to ten minutes before cutting.
3. **Float sections in a 40 °C water bath.** Fill a flotation water bath with distilled water and heat it exactly to 40 °C. Use fine forceps or a paintbrush to gently place the cut ribbon (or individual sections) on the water surface, with the shiny side (the original block face) facing down. Let the sections flatten completely, which usually takes five to fifteen seconds; avoid floating them for more than one minute, as this can cause the tissue to expand or become disrupted. If needed, a brush can be used to carefully separate individual sections.
4. **Collect sections onto glass slides.** Use clean, positively charged glass slides to improve tissue adherence. Gently dip the slide into the water bath at a slight angle, catching one section per slide or multiple sections per slide as desired. Lift the slide vertically to pick up the section, taking care not to trap air bubbles beneath it. For serial sections, collect consecutive sections onto separate slides or arrange them in order on a single slide.
5. **Dry slides at room temperature or at 40 °C.** For room-temperature drying, leave them overnight (12 to 24 hours) in a dust-free environment. For drying at 40 °C, place the slides on a slide warmer or in a forced-air incubator for one to two hours. Ensure sections are completely dry before staining or storage, as incomplete drying may cause them to detach during subsequent staining steps.

**Figure 7.**
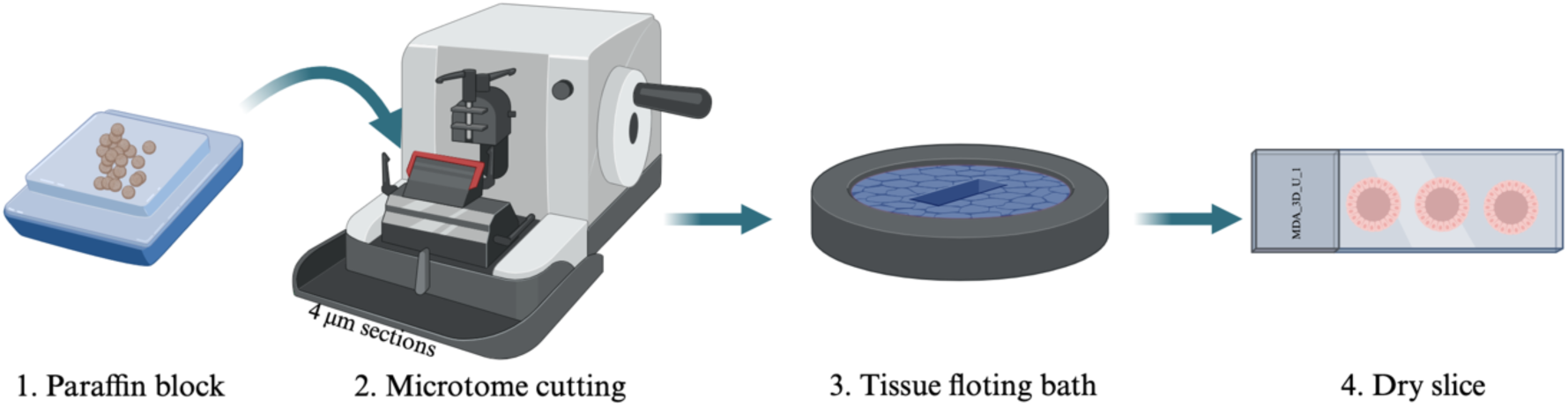
Microtome sectioning of paraffin-embedded spheroids.

**Note:** For small spheroids, carefully trim the block to avoid cutting too deeply past the sample. Clean the water bath daily to prevent contamination. Store dried slides in a closed slide box at room temperature, protected from dust and moisture. If sections wrinkle repeatedly, add a drop of 0.1% gelatin or a commercial adhesive to the water bath.

### X. Hematoxylin and eosin (H&E) staining (**Figure 8**)

After sectioning and drying, slides must be stained to visualize nuclear and cytoplasmic details. Perform all steps at room temperature in a fume hood when using xylene. Use fresh solutions and change them according to the laboratory schedule.

1. Heat the slides at 100 °C for 5 minutes to melt the paraffin and firmly adhere the sections to the glass. Place the slides on a slide warmer or in a preheated oven, ensuring they are level to prevent sections from running off.
2. Immerse the slides in xylene three times for 5 minutes each. Xylene removes all remaining paraffin. Use a clean xylene bath for each immersion, and gently agitate the slides periodically to ensure effective deparaffinization.
3. Transfer the slides to 96% ethanol for three 3-minute changes. This step gradually replaces xylene with alcohol, rehydrating the tissue.
4. Immerse the slides in 80% ethanol for 3 minutes to continue the graded rehydration.
5. Immerse the slides in 70% ethanol for 3 minutes to bring the sections to a lower alcohol concentration.
6. Rinse the slides briefly in distilled water to remove ethanol and prepare the tissue for aqueous staining solutions. Use a gentle stream or a dip in a clean water bath.
7. Immerse the slides in hematoxylin solution for 7 minutes. Hematoxylin stains cell nuclei a deep blue-purple. Use a standard hematoxylin. Ensure the slides are fully submerged and not touching each other.
8. Rinse the slides in distilled water to remove excess hematoxylin. A single gentle rinse is usually sufficient; avoid over-rinsing, which may wash out the stain.
9. Dip the slides in acid alcohol with three quick immersions. Differentiation removes excess hematoxylin from the cytoplasm and connective tissue, leaving nuclei sharply stained. Do not leave slides in acid alcohol for more than a few seconds.
10. Immerse the slides in lithium carbonate solution (a weak alkaline solution, usually 0.1-0.5% lithium carbonate in water) for a few seconds. This step “blues” the hematoxylin, converting it from a red-brown to a blue-purple color.
11. Rinse the slides in distilled water to remove residual lithium carbonate.
12. Immerse the slides in 70% ethanol for 1 minute to re-equilibrate the tissue before eosin staining.
13. Immerse the slides in eosin solution for 10 minutes. Eosin stains cytoplasmic components pink to orange. Use an alcoholic or aqueous eosin as preferred.
14. Transfer the slides to 80% ethanol for 3 minutes to begin removing water and excess eosin.
15. Immerse the slides in 96% ethanol for three changes of 3 minutes each. Complete dehydration is essential for proper clearing.
16. Immerse the slides in xylene for two changes of 5 minutes each. Xylene replaces ethanol, making the sections transparent and ready for mounting.
17. Air-dry the slides in a fume hood for 5-10 minutes, or until no traces of xylene remain. Do not let the sections become completely dry before mounting, as this can cause cracking. Alternatively, mount immediately from the last xylene bath using a resinous mounting medium.

**Figure 8.**
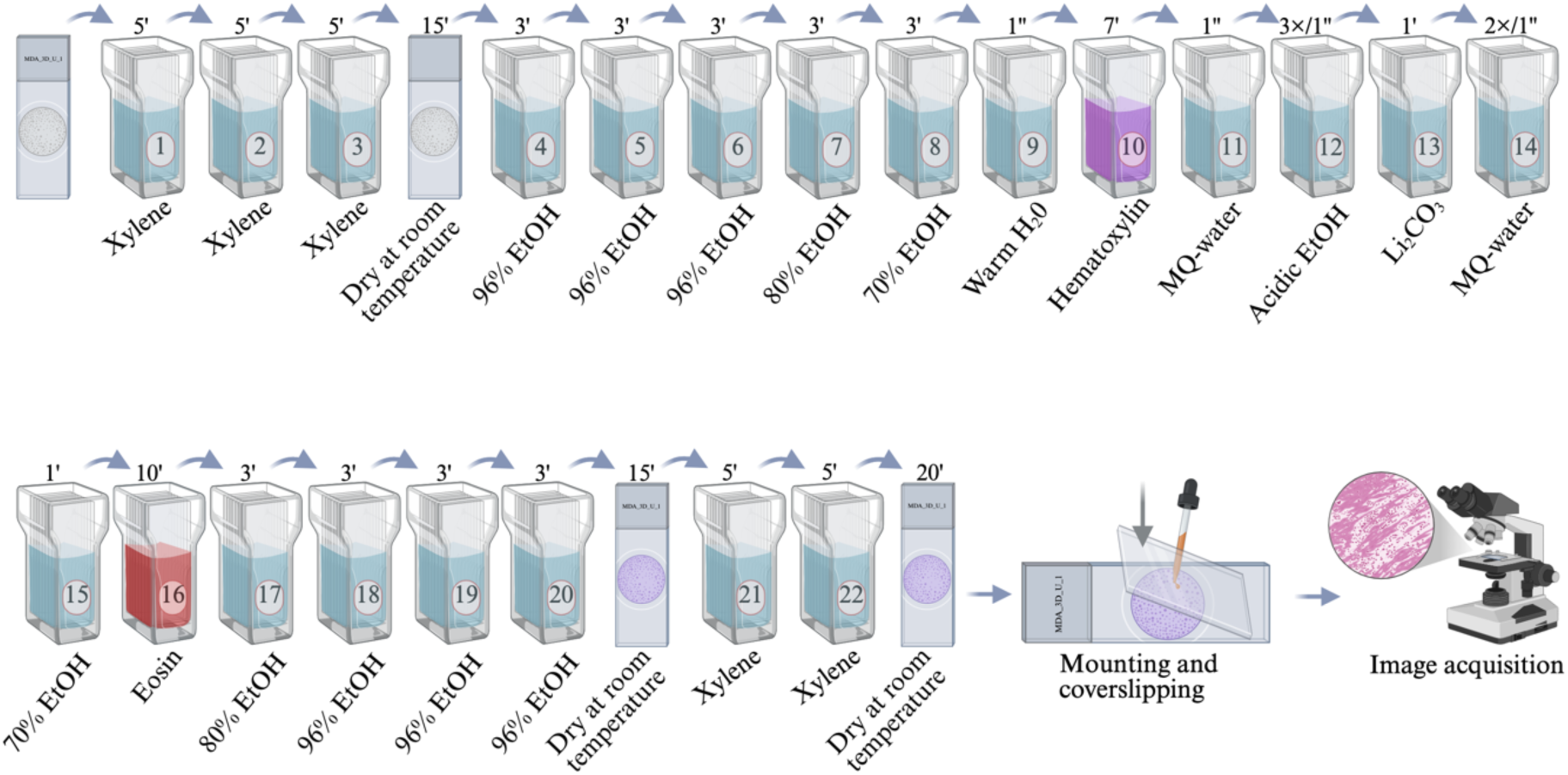
Spheroid sections stained with hematoxylin and eosin (H&E) and coverslipped.

**Note:** Always wear appropriate personal protective equipment (gloves, lab coat, safety glasses) when handling xylene, acid alcohol, and lithium carbonate. If background staining is too dark, shorten the eosin step or extend differentiation. Store stained slides in a dust-free slide box after coverslipping.

### XI. Mounting and coverslipping of stained sections (**Figure 8**)

After clearing, slides must be coverslipped with a resinous mounting medium to preserve the stained sections for long-term storage and microscopic examination. Perform this step immediately after the final xylene bath to prevent the sections from drying out and crystallizing.

1. Apply mounting resin immediately after the last xylene step. Do not let the slides air-dry completely, as dried sections become brittle and hard to cover without air bubbles. Remove the slide from xylene, drain excess solution by touching the edge to a paper towel, and work quickly.
2. Add one drop of mounting resin directly onto the section. Use a clear, xylene-based resin. The drop should be just large enough to spread under the coverslip without overflowing. Avoid using too much resin, which can seep out and become sticky.
3. Place a clean glass coverslip carefully over the drop. Lower the coverslip at a slight angle (approximately 45 degrees) using fine forceps or a needle, allowing the resin to spread evenly without trapping air bubbles (Figure 8). Gently press the coverslip down with the back of the forceps to expel any small bubbles and ensure uniform coverage.
4. Allow the mounted slides to dry at room temperature in a horizontal position for 24-48 hours. Protect the slides from dust during drying by placing them in a covered slide box or under a dust cover. Once fully hardened, slides can be stored upright in a slide box at room temperature.

**Note:** Always work in a fume hood when using xylene-based mounting resins. Clean any excess resin from the edges of the coverslip with a lint-free tissue moistened with xylene. For faster drying, slides can be placed on a slide warmer at 37 °C for 2-4 hours, but room temperature curing results in a more durable mount.

### XII. Image acquisition

After mounting and drying, slides should be examined and photographed to record staining results. Use an inverted microscope equipped with a digital camera and suitable brightfield optics. Conduct all imaging under consistent lighting and magnification for each experimental condition.

1. Acquire images with an inverted microscope (Figure 8). Place the slide on the stage with the coverslip facing the objective lens. Begin with low magnification (e.g., 4× or 10×) to locate spheroids or regions of interest, then switch to higher magnification (e.g., 20× or 40×) for detailed observation (Figures 9 and 10). Adjust the focus, brightness, and contrast to produce a clear, evenly illuminated image. Use the same exposure time and white balance settings for all samples to ensure consistency.
2. Capture representative images for each experimental condition. For each condition (such as control, treated, or various time points), take at least three to five images from non-overlapping fields. Include images at low magnification to display overall spheroid morphology and at high magnification to detail nuclear and cytoplasmic staining. Save images in an uncompressed format (like TIFF) with a consistent naming convention that includes the date, sample identifier, condition, and magnification.

**Figure 9.**
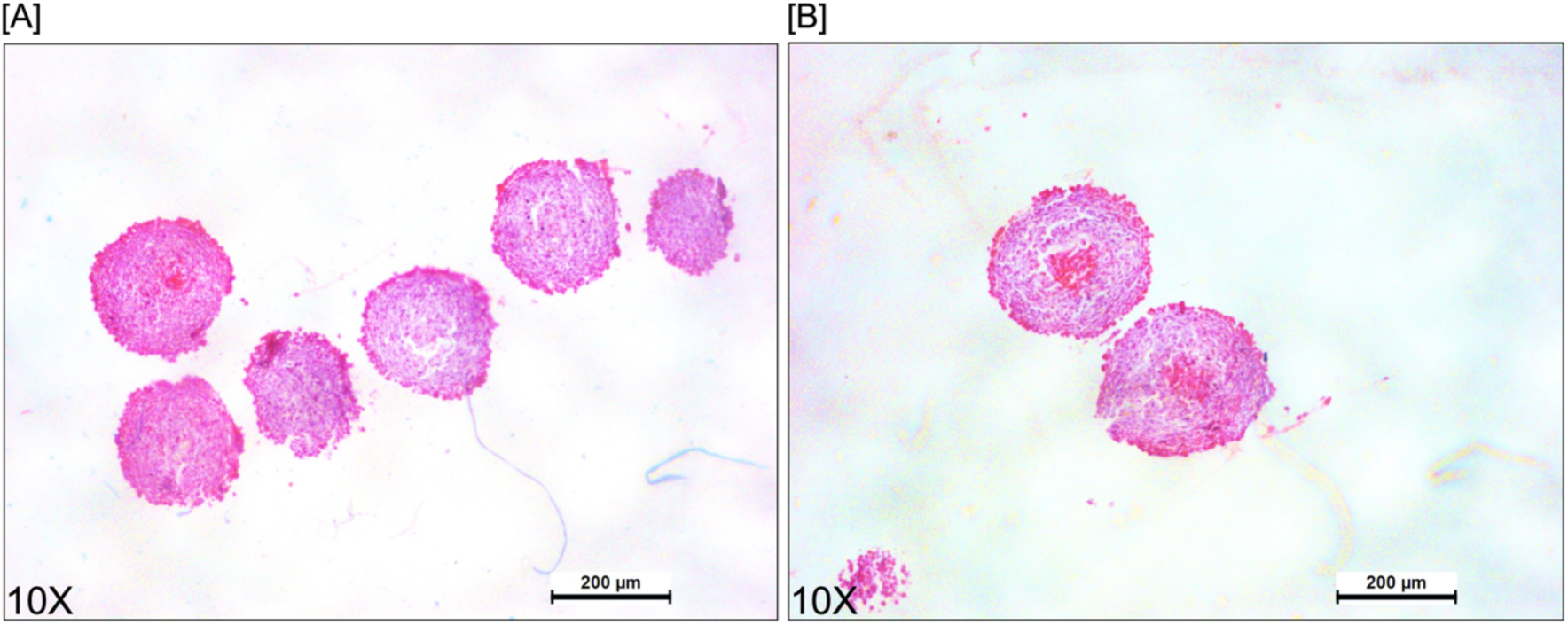
Histological analysis of MDA-MB-231 spheroids stained with hematoxylin and eosin (H&E). (A) Representative serial sections from a single spheroid, showing internal structural consistency. (B) 10X magnification view highlighting the characteristic zonal architecture of a mature spheroid, clearly delineating the peripheral proliferative zone, the intermediate quiescent region, and the central necrotic core.

**Figure 10.**
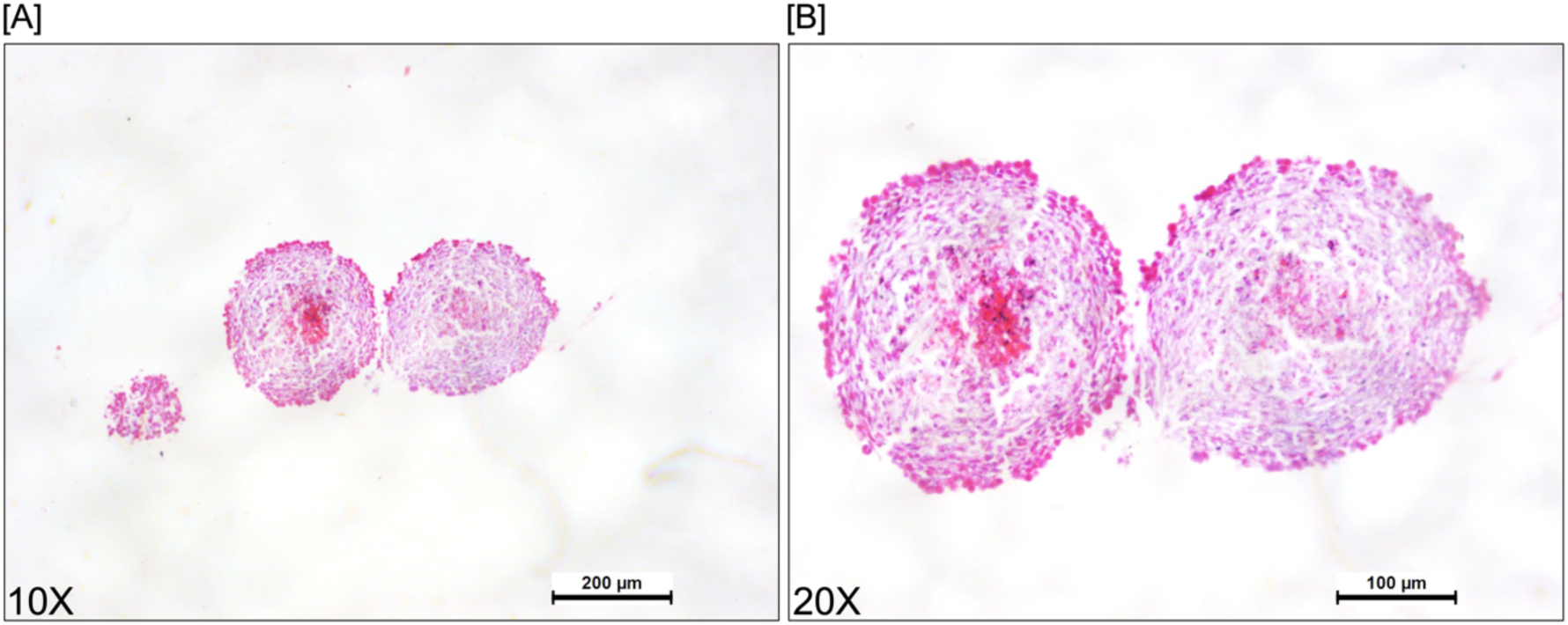
Sections of a mature MDA-MB-231 spheroid stained with hematoxylin and eosin (H&E). (A) Representative sections showing the typical spheroid structure (10X magnification). (B) A 20X magnification view emphasizing the distinctive zonal architecture of a mature spheroid, clearly illustrating the outer proliferative zone, the intermediate quiescent region, and the central necrotic core.

**Note:** Calibrate the microscope with a stage micrometer before acquisition if you plan to perform quantitative measurements like spheroid diameter or staining intensity. Avoid overexposure to prevent bleaching eosin signals, and use a clean objective lens for each session. Organize image files in a structured folder system and keep a backup copy.

### Validation of protocol

#### Morphological and histological evaluation of Actinomycin D efficacy in MDA-MB-231 spheroids

Spheroids derived from the MDA-MB-231 cell line are a well-established, physiologically relevant preclinical model in cancer research. This cell line originated from a triple-negative breast cancer (TNBC) patient, characterized by the absence of estrogen receptor (ER), progesterone receptor (PR), and HER2 amplification, making it inherently resistant to conventional hormone-and HER2-targeted therapies. As a result, MDA-MB-231 spheroids serve as a valuable platform for testing new anti-cancer compounds and understanding their mechanisms of action in conditions that more closely mimic the tumor microenvironment than standard monolayer cultures.

The morphological assessment of compound-treated spheroids serves two main purposes: it offers an initial indication of cytotoxic effectiveness while also providing indirect evidence of drug penetration into the three-dimensional (3D) architecture. A compound that does not diffuse beyond the outer cell layers usually causes surface damage without affecting the core, whereas an effective penetrant compound induces structural changes throughout the entire spheroid. In this context, treatment with Actinomycin D, a powerful transcription inhibitor and well-known chemotherapeutic agent, produced clear phenotypic changes in MDA-MB-231 spheroids. Treated cultures showed a significant and consistent reduction in overall spheroid diameter, along with a gradual loss of the typical compact, round morphology (Figure 11). Additionally, the appearance of many rounded, highly refractile (phase-bright) bodies at the spheroid edges and in the surrounding medium-morphologically similar to apoptotic cells-provided clear visual evidence of extensive programmed cell death. Taken together, these observations confirm that Actinomycin D effectively penetrates the 3D structure and exerts strong cytotoxic activity against this aggressive triple-negative breast cancer (TNBC) model, supporting its use as a positive control or reference compound in future spheroid experiments.

**Figure 11.**
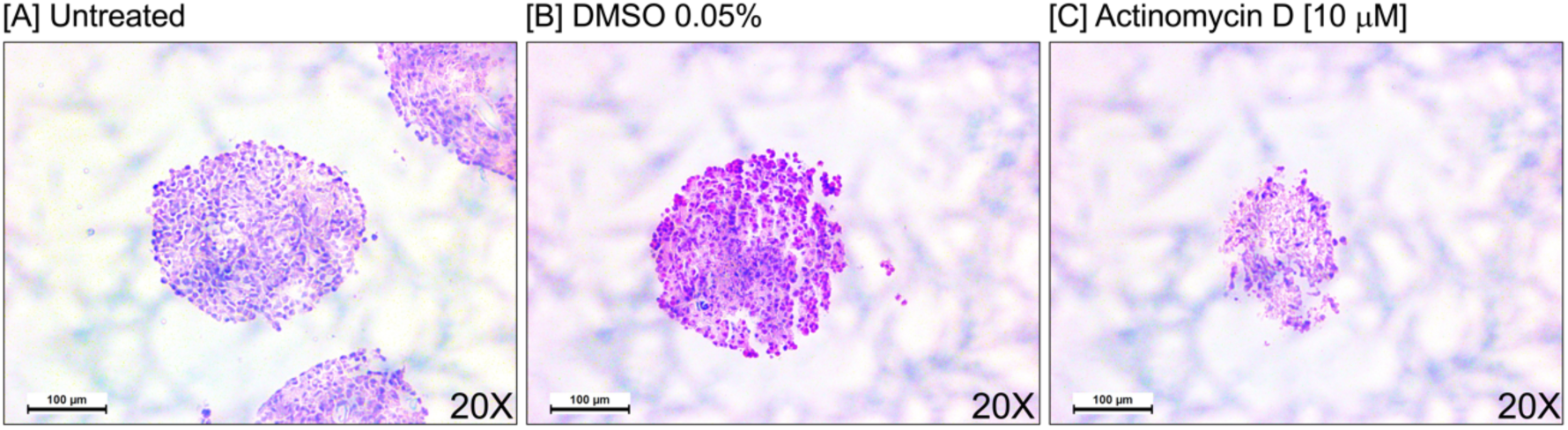
Morphological effects of Actinomycin D on MDA-MB-231 spheroids. Representative bright-field images of spheroids under three conditions: (A) Untreated control, showing typical compact and rounded morphology; (B) Vehicle control (DMSO 0.05%), confirming no solvent-induced effects on spheroid integrity; and (C) Actinomycin D (10 µM) for 48 h, showing noticeable changes in spheroid size and structural organization.

#### Histological evaluation of Actinomycin D-induced cytotoxicity in A-549 spheroids

The A-549 cell line, derived from human lung adenocarcinoma, is one of the most widely used models in non-small cell lung cancer research. Its extensive use is due to its well-documented genetic background and predictable behavior in both 2D and 3D culture systems. When grown under non-adherent conditions, A-549 cells naturally form cohesive, relatively compact spheroids with well-defined, smooth borders and consistent size distribution. This high level of reproducibility makes A-549 spheroids an excellent platform for high-throughput drug screening, allowing for reliable quantitative assessments of compound effectiveness across multiple experimental runs.

Morphological evaluation of treated spheroids provides critical insights that extend beyond simple cytotoxicity. Specifically, the degree and pattern of structural disruption serve as a surrogate marker for drug penetration capacity: compounds that fail to reach the inner layers typically cause surface erosion without compromising the core, whereas agents with favorable diffusivity induce global architectural destabilization. Exposure of A-549 spheroids to Actinomycin D produced striking and concentration-dependent phenotypic changes (Figure 12). Treated cultures displayed a substantial and progressive reduction in spheroid diameter, accompanied by progressive loss of surface smoothness and overall structural compaction.

**Figure 12.**
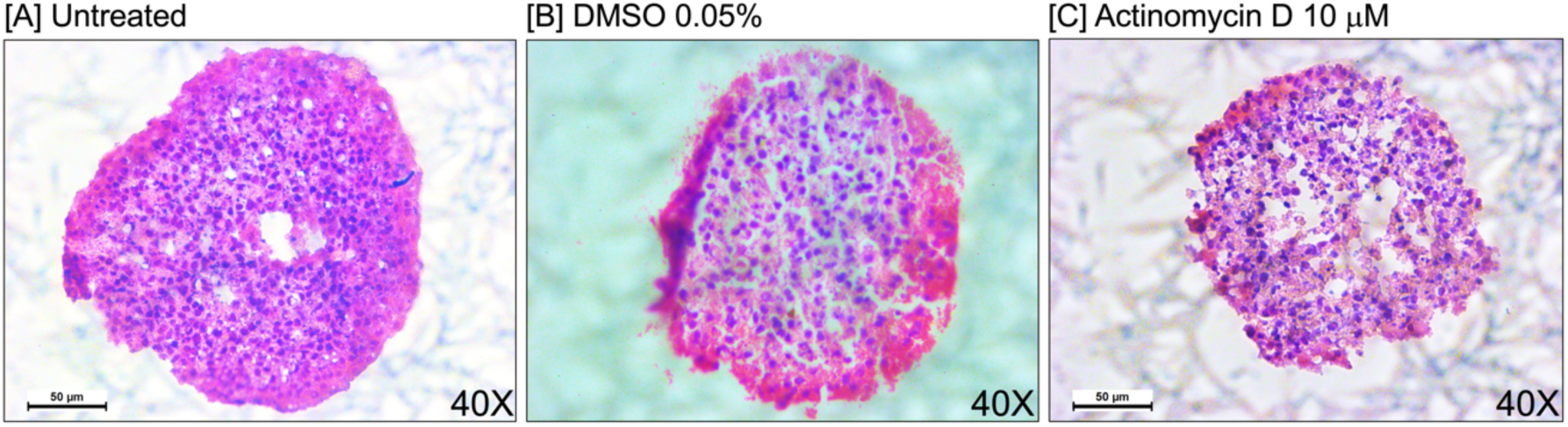
Actinomycin D disrupts A549 spheroid morphology and induces apoptosis. Representative bright-field images of A549 lung carcinoma spheroids under three experimental conditions: (A) Untreated control, showing a dense, compact, and well-rounded shape with smooth borders; (B) Vehicle control (DMSO 0.05%), confirming that the solvent does not affect spheroid integrity or structure; and (C) Actinomycin D (10 µM). Treated spheroids display significant structural breakdown, including decreased size, loss of cohesion, surface irregularities, and extensive cellular detachment from the spheroid edges.

#### Visualizing Actinomycin D-induced structural alterations in CaCo-2 spheroids through histology

The CaCo-2 cell line, originally derived from a human colorectal adenocarcinoma, is widely used in pharmaceutical research, especially for studying intestinal permeability and drug transport mechanisms. However, when cultured under non-adherent conditions to form 3D spheroids, CaCo-2 cells pose a distinct morphological challenge: unlike the compact, uniformly rounded aggregates seen in breast or lung carcinoma models, CaCo-2 cells naturally form looser, irregularly shaped, and structurally diverse assemblies with poorly defined borders and variable cell-cell adhesion. This inherent fragility makes whole-mount brightfield assessment difficult, as baseline morphological variability can hide subtle treatment-related changes. Therefore, establishing a robust histological pipeline—including careful retrieval, formalin fixation, paraffin embedding, and serial H&E sectioning—is especially important for this model. Sectioning enables standardized, high-resolution visualization of internal cytoarchitecture, effectively overcoming limitations imposed by surface-level heterogeneity.

At the 48-hour mark, Actinomycin D treatment (10 µM) already causes easily detectable cytopathic effects that exceed the natural variability of untreated cultures. Histological examination of control spheroids, both untreated and vehicle-treated, shows loosely cohesive cell clusters with scattered intercellular spaces, consistent with the characteristic phenotype of this line. In sharp contrast, samples exposed to Actinomycin D display significant architectural erosion, marked by widespread cellular dissociation, loss of intercellular contacts, and the development of large acellular zones within the aggregate matrix (Figure 13). Additionally, H&E-stained sections prominently feature numerous condensed, hyperchromatic nuclei alongside fragmented eosinophilic cytoplasmic remnants, indicating active apoptosis, throughout the remaining cellular mass. Overall, this histological analysis confirms the compound’s strong anti-tumor activity even in challenging, less compact 3D structures, while also showing that CaCo-2 spheroids are suitable for screening drugs targeting colorectal cancers, as long as appropriate histological endpoints are used.

**Figure 13.**
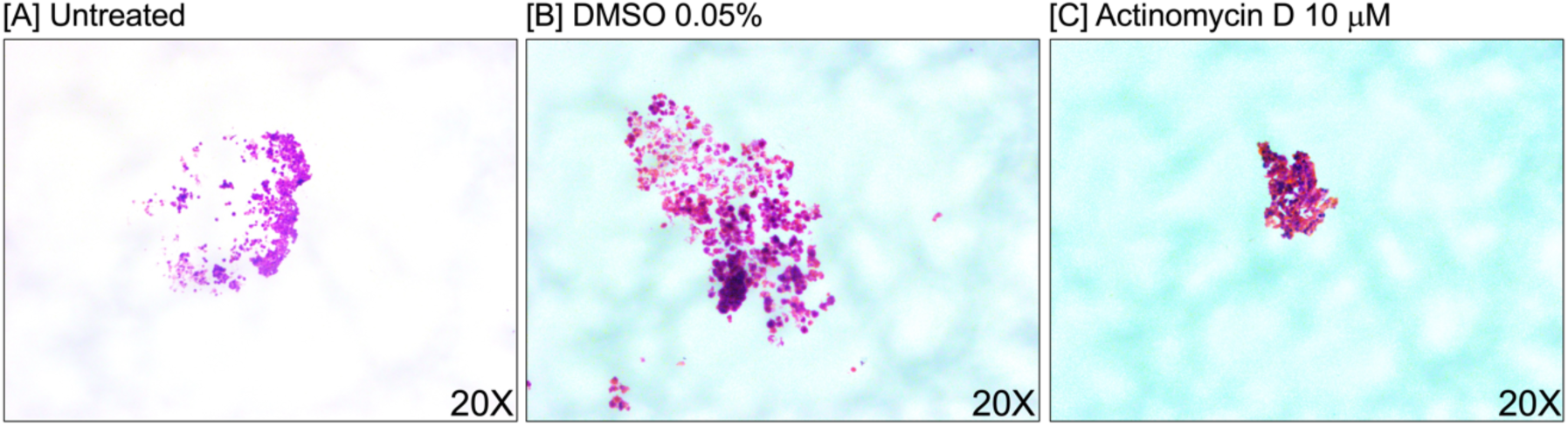
Actinomycin D causes cytotoxic disassembly of CaCo-2 spheroids. Phase-contrast micrographs of CaCo-2 colorectal spheroids taken 48 h after treatment. (A) Untreated cultures and (B) vehicle controls (DMSO 0.05%) both show the characteristic loose, granular structure typical of this cell line, with minimal background cellular debris. (C) The addition of Actinomycin D (10 µM) induces a clear phenotypic change: the spheroid matrix visibly collapses, along with extensive shedding of membrane-blebbed remnants and highly condensed apoptotic figures. Notably, despite the naturally limited compaction of CaCo-2 spheroids, the cytotoxic signature remains unequivocally distinguishable from the basal state.

#### General notes and troubleshooting

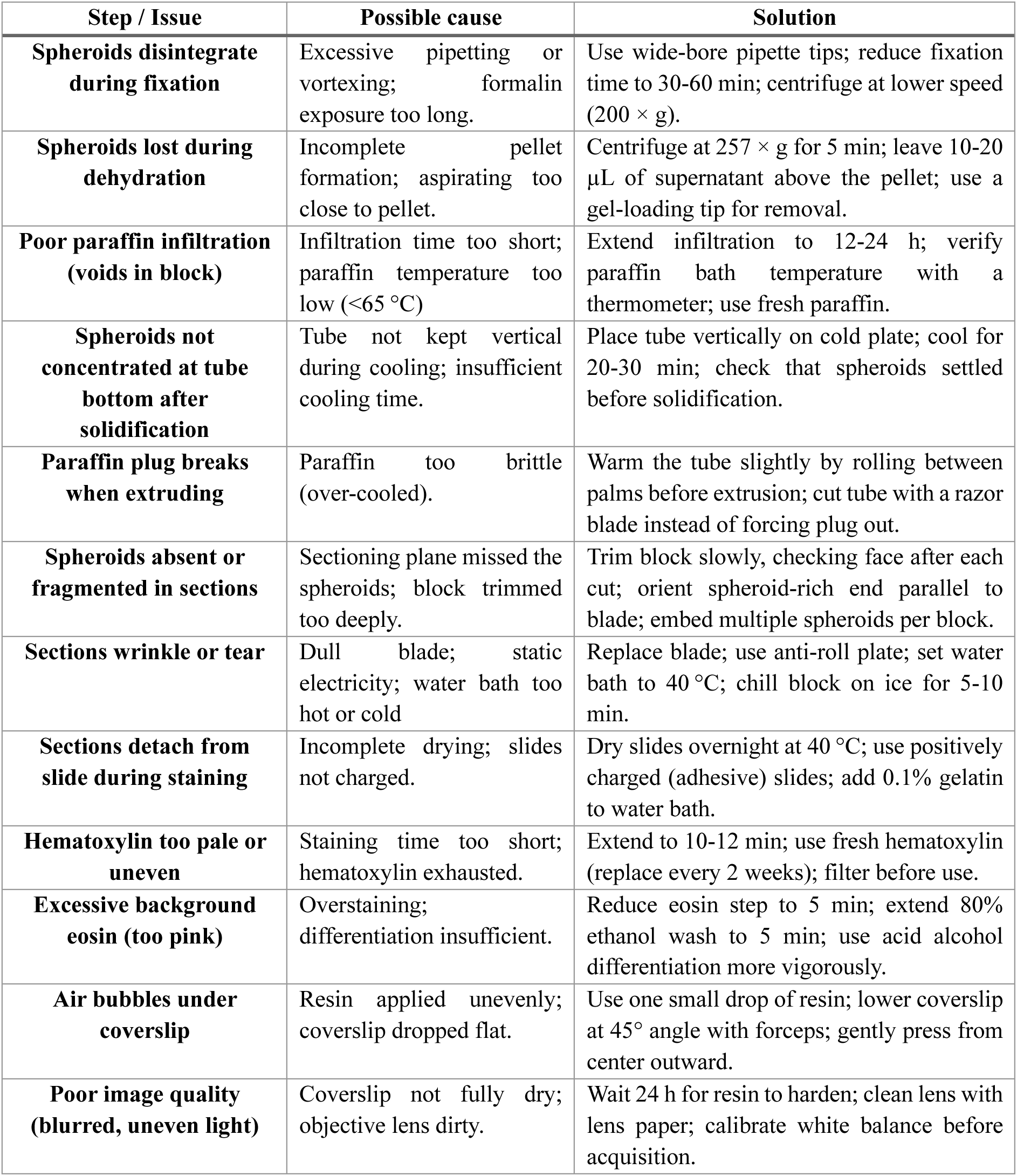

## Acknowledgment

We gratefully acknowledge funding from the Coordinación de la Investigación Científica de la UMSNH (Proyecto 17644). Ramón Cervantes-Rivera thanks the Secretaría de Ciencia, Humanidades, Tecnología e Innovación (SECIHTI) for postdoctoral stipend support.

## Author contribution

**Conceptualization:** Ramón Cervantes-Rivera

**Methodology:** Ramón Cervantes-Rivera, Adrián Sánchez Orozco, Ma. Antonia Herrera-Vargas

**Investigation:** Ramón Cervantes-Rivera, Atalia Ziret Romero Rosas, Sandra Jetsamari Figueroa Ortíz

**Formal analysis:** Ramón Cervantes-Rivera, Atalia Ziret Romero Rosas, Sandra Jetsamari Figueroa Ortíz

**Data curation:** Ramón Cervantes-Rivera

**Writing-Original draft:** Ramón Cervantes-Rivera, Sandra Jetsamari Figueroa Ortíz

**Writing-Review & Editing:** Ramón Cervantes-Rivera, Atalia Ziret Romero Rosas, Alejandra Ochoa Zarzosa, Joel E. López-Meza, Esperanza Meléndez-Herrera and Manuel López Rodríguez

**Visualization:** Ramón Cervantes-Rivera, Sandra Jetsamari Figueroa Ortíz

**Funding acquisition:** Joel E. López-Meza and Alejandra Ochoa Zarzosa

**Supervision:** Ramón Cervantes-Rivera, Alejandra Ochoa Zarzosa and Joel E. López-Meza

**Project administration:** Ramón Cervantes-Rivera and Joel E. López-Meza

## Competing interests

The authors declare no conflict of interest.

## Notes

### Competing Interest Statement

The authors have declared no competing interest.

